# Identification and characterisation of a phospholipid scramblase in the malaria parasite *Plasmodium falciparum*

**DOI:** 10.1101/2020.06.22.165258

**Authors:** Silvia Haase, Melanie Condron, David Miller, Dounia Cherkaoui, Sarah Jordan, Jacqueline M Gulbis, Jake Baum

## Abstract

Recent studies highlight the emerging role of lipids as important messengers in malaria parasite biology. In an attempt to identify interacting proteins and regulators of these dynamic and versatile molecules, we hypothesised the involvement of phospholipid translocases and their substrates in the infection of the host erythrocyte by the malaria parasite *Plasmodium* spp. Here, using a data base mining approach, we have identified a putative phospholipid (PL) scramblase in *P. falciparum* (*Pf*PLSCR) that is conserved across the genus and in closely related unicellular algae. By reconstituting recombinant *Pf*PLSCR into liposomes, we demonstrate metal ion dependent PL translocase activity and substrate preference, confirming *Pf*PLSCR as a *bona fide* scramblase. We confirm that *Pf*PLSCR is expressed during asexual and sexual parasite development, localising to different membranous compartments of the parasite throughout the intra-erythrocytic life cycle. Two different gene knockout approaches, however, suggest that *Pf*PLSCR is not essential for erythrocyte invasion and asexual parasite development, pointing towards a possible role in other stages of the parasite life cycle.

## Introduction

Infection of the host erythrocyte by the malaria parasite *Plasmodium* spp. is a crucial event in the pathobiology of this pathogen. The process of erythrocyte invasion involves a highly orchestrated cascade of protein-protein interactions between the parasite and the host cell. Once inside, the asexual, intra-erythrocytic proliferation of *Plasmodium* spp. is accompanied by a high demand of lipids. These lipids are required to build membranes for daughter cell and organelle neogenesis, promote the expansion of the parasitophorous vacuole (PV) in which the parasite resides and develops, and to generate a parasite-derived exomembrane system involved in parasite virulence and nutrient uptake [reviewed in (1,2)]. Accordingly, the progression of blood-stage parasites from ring to schizont stages is marked by increased metabolic activity and stage-specific changes in the lipidomic profiles within the infected erythrocyte (3). Individual phospholipids have also come into focus by playing fundamental roles in signalling. Phosphoinositides have been implicated in merozoite development (4), haemoglobin digestion (5), microneme secretion (6) and apicoplast biogenesis in *Plasmodium* spp. and its apicomplexan relative *Toxoplasma gondii* (7,8). Another important lipid messenger, phosphatidic acid, has also been linked to microneme exocytosis (9) and host cell egress in *Toxoplasma gondii* (10). Both, the apicoplast (a non-photosynthetic, relict plastid shared by most members of the phylum Apicomplexa) and the micronemes (specialised organelles containing adhesins involved in erythrocyte invasion) are essential to parasite survival. Lipid rafts have also been implicated in facilitating and regulating host-pathogen interactions in different protozoan parasites, including *Plasmodium* spp. and the intestinal pathogens *Giardia intestinales* and *Entamoeba histolytica* [reviewed in (11)]. Investigations into the roles played by lipids and their interacting proteins and regulators are therefore increasingly seen as an attractive avenue to explore and to identify novel antimalarial and antiparasitic drug targets.

Prompted by the growing evidence and importance of lipid function in malaria parasite and apicomplexan biology, we sought to investigate the role of putative phospholipid (PL) translocases and their substrates in the infection of the host erythrocyte by the most virulent human malaria parasite *P. falciparum*, which causes an estimated ~ 400,000 fatalities and ~ 200 million cases per year worldwide (WHO report 2019, www.who.int). An initial search for candidates in the *Plasmodium* Genomics Resource (www.plasmodb.org) revealed a predicted PL scramblase which fulfilled all characteristics of our search rationale: (i) putative PL translocase activity, (ii) maximal expression in the late blood stages and (iii) predicted localisation to the plasma membrane in the invasive stage, the merozoite.

Phospholipids are asymmetrically distributed in the plasma membrane. While phosphatidylcholine (PC) and sphingomyelin (SM) dominate the outer leaflet of the lipid bilayer, phosphatidylserine (PS), phosphatidylethanolamine (PE) and phosphatidyl-inositol (PI) characterise the inner leaflet. Phospholipid asymmetry is maintained by PL translocases, such as flippases and floppases, which function as ATP-dependent unidirectional transporters that move PLs from the outer to the cytosolic face or from the inner to the exoplasmic face, respectively [reviewed in (12)]. In contrast, PL scramblases, including members of the PLSCR, TMEM16 and XKR families, are activated by an increase of intracellular Ca^2+^ in response to cell stress or extracellular signals such as apoptotic stimuli and mediate the bidirectional movement of PLs across the bilayer. Consequently, PS and other lipids normally confined to the inner leaflet become exposed at the cell surface and aid in the process of blood coagulation and clearance of dead or injured cells [reviewed in (13)].

The human genome encodes four members of the PLSCR family (hPLSCR1-4); with the exception of hPLSCR2, these proteins have been reported to be expressed in various tissues (14). The best studied member to date is hPLSCR1; initially identified in and purified from erythrocytes (15,16), the protein is characterised by several functional domains: an N-terminal proline-rich domain containing multiple PXXP and PPXY motifs suggested to mediate protein-protein interactions (14); a DNA binding motif spanning Met^86^ to Glu^118^ which has been implicated in binding the promoter region of the inositol 1,4,5-triphosphate receptor type I (17); a palmitoylation site ^184^CCCPCC^189^ involved in trafficking to the plasma membrane (18); a non-classical nuclear leader sequence (NLS) ^257^GKISKHWTGI^266^ shown to bind importin α (19) precedes an EF-hand like Ca^2+^ binding motif ^273^D[A/S]DNFGIQFPLD^284^ which is directly followed by a C-terminal α-helical stretch suggested to insert hPLSCR1 into artificial bilayers (16,20,21). The integral membrane protein is expressed in blood cells and a wide range of tissues and cancer cell lines (15,16,22) and is capable of translocating PLs in reconstituted membrane systems (15,20,23–25) While its actual cellular role in PL scrambling remains a matter of debate, hPLSCR1 has been implicated in a range of different activities related to apoptosis, transcriptional regulation, autoimmunity, antiviral defence and the development of cancer and lipid-related diseases [reviewed in (13)].

In this study, we biochemically characterised recombinant *Pf*PLSCR and applied a gene tagging approach to investigate the expression and localisation of endogenously expressed *Pf*PLSCR-HA in transgenic *P. falciparum* parasites. In addition, we have utilised two different gene knockout strategies to analyse the loss of function phenotype in mutant and *Pf*PLSCR deficient parasite lines. Though the function of the protein remains elusive, our findings support the biochemical and cellular observations made for the human ortholog hPLSCR1 and may suggest a still to be determined regulatory role in parasite sexual development.

## Results

### *P. falciparum* encodes a putative PL scramblase

In order to identify putative PL translocases, we used three search criteria available in the *Plasmodium* Genomics Resource database (www.plasmodb.org): (i) Function prediction as determined by EC number and GO term, (ii) transcriptomics data and (iii) protein features and properties. This search identified a single exon gene (gene identifier PF3D7_1022700) encoding for a putative PL scramblase of approx. 33 kDa, hereafter referred to as *Pf*PLSCR (Fig. 1A), which is predicted to be highly upregulated in late schizont stages (40-48h post invasion) and to associate with the plasma membrane.

**Figure 1.**
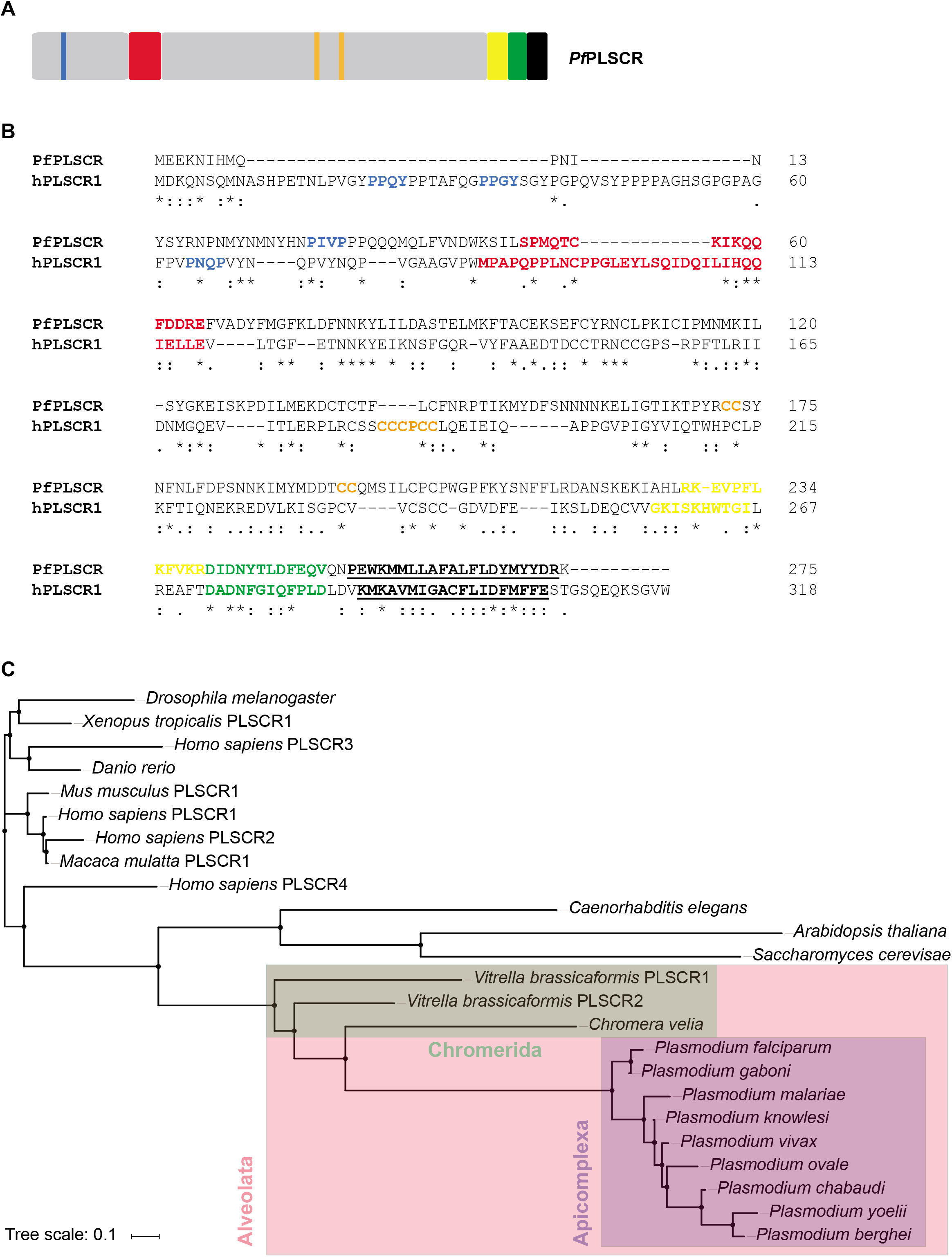
Protein features and conservation of the putative PL scramblase in *P. falciparum*. A) A schematic of *Pf*PLSCR indicating the PXXP/Y motifs (blue), putative DNA binding region (red), predicted palmitoylation sites (orange) and bipartite nuclear leader sequence (yellow), as well as a putative Ca2+ binding motif (green) and the C-terminal transmembrane helix (black and underlined). B) Alignment of *Pf*PLSCR (PlasmoDB ID: PF3D7_1022700) with the human ortholog hPLSCR1 (UniProtKB ID: O15162). The functional domains, colour-coded as above, are positionally conserved in *Pf*PLSCR. Similar residues are marked by colons (:) and identical residues are indicated by stars (*). Protein sequences were aligned with Clustal Omega (67). The cut-off score was set to 4.0 for the prediction of palmitoylation sites (68). C) Phylogenetic tree reconstruction from a selection of PLSCR orthologues. The tree was generated by using the Simple Phylogeny Web services (69) and the tree file was imported into iTOL (70) for illustrative purposes. The following EupathDB gene identifiers were used: PF3D7_1022700 (*P. falciparum*), PGABG01_1020700 (*P. gaboni*), PmUG01_10033900 (*P. malariae*), PKH_060680 (*P. knowlesi*), PVX_111580 (*P. vivax*), PocGH01_06015800 (*P. ovale*), PCHAS_050700 (*P. chabaudi*), PY17X_0508000 (*P. yoelii*), PBANKA_050690 (*P. berghei*), Vbra_21596 (*Vitrella brassicaformis* PLSCR1), Vbra_2039 (*Vitrella brassicaformis* PLSCR2) and Cvel_12647 (*Chromera velia*). The following UniProtKB accession numbers were used: O15162 (*H. sapiens* PLSCR1), Q9NRY7 (*H. sapiens* PLSCR2), Q9NRY6 (*H. sapiens* PLSCR3), Q9NRQ2 (*H. sapiens* PLSCR4), Q9JJ00 (*Mus musculus* PLSCR1), H9ZA71 (*Macaca mulatta* PLSCR1), Q21318 (*Caenorhabditis elegans*), P47140 (*Saccharomyces cerevisae*), Q8IQD8 (*Drosophila melanogaster*), Q6NY24 (*Danio rerio*) and B5DDV8 (*Xenopus tropicalis* PLSCR1).

A protein sequence alignment of *Pf*PLSCR with hPLSCR1 revealed moderate sequence conservation between the two orthologs sharing only 17% sequence identity and 31% sequence similarity [www.bioinformatics.org/sms2/ident_sim.html (26)]. However, all characteristic domains of hPLSCR1 appear to be partially or fully conserved in the parasite ortholog (Fig. 1B). The N-terminus of *Pf*PLSCR is shorter but contains several proline residues with a possible PXXP motif. A putative, semi-conserved DNA binding region Ser^50^ to Glu^65^ precedes two predicted palmitoylation sites ^172^CC^173^ and ^194^CC^195^ [http://csspalm.biocuckoo.org (27)] and a predicted bipartite NLS which starts at Arg^228^ [http://nls-mapper.iab.keio.ac.jp (28)] and partially overlaps with the putative Ca^2+^ binding site. Several residues of the Ca^2+^ binding region ^240^DIDNYTLDFEQV^251^, most notably in positions 1 (Asp^240^), 3 (Asp^242^) and 9 (Phe^248^), as well as its positioning adjacent to the C-terminal transmembrane helix spanning Pro^254^ to Arg^274^ [http://topcons.cbr.su.se (29)] are conserved in the putative PL scramblase of *P. falciparum*, indicating a functional preservation of these domains in the parasite protein.

### *Pf*PLSCR is conserved within *Plasmodium* spp. but not in other Apicomplexa

Orthologs of hPLSCR1 are present in various organisms ranging from fungi and plants to animals [reviewed in (30)]. A BLAST analysis against the Eukaryotic Pathogen database (www.eupathdb.org) revealed putative PLSCR proteins across the genus *Plasmodium*. Protein sequence identities to *Pf*PLSCR range from approx. 68% for the orthologs of the rodent malaria parasites *P. berghei and P. yoelii* to 97% for the putative PLSCR of *P. gaboni*, a chimpanzee infecting member of the *P. falciparum* lineage (Fig. 1C). While the N-termini of the *Plasmodium* spp. PLSCR proteins displayed some variation in length and in the abundance of proline residues, the functional domains of the C-terminal portion appeared to be highly conserved as previously described for other PLSCR orthologs [reviewed in (30)]. No putative PLSCR proteins could be identified in the remaining Apicomplexa or unicellular pathogens, including members of the phyla Amoebozoa, Diplomonadida, Kinetoplastida, Oomycetes and Trichomonanida, pointing towards a specific role of the PL scramblase in malaria parasite biology.

The search, however, did identify orthologs in the photosynthetic algae *Chromera velia* and *Vitrella brassicaformis*, sharing 24-28% sequence identity and 37-42% sequence similarity with the putative PL scramblase of *P. falciparum* (Fig. S1). The algae represent the closest known relative of apicomplexan parasites and are classified as Chromerida, a marine phylum belonging to an ancient group of unicellular eukaryotes, the Alveolata, which are characterised by flattened, membranous sacs (alveoli) just underneath the plasma membrane (31–33).

### Overexpression and purification of recombinant *Pf*PLSCR

To biochemically characterise the putative *Pf*PLSCR and validate its potential PL translocase activity, we recombinantly expressed the full-length protein with a N-terminal hexa-histidine affinity tag (His::*Pf*PLSCR) in *E. coli* BL21(DE3). Several attempts to obtain biologically active His::*Pf*PLSCR using previously described methods for recombinant hPLSCR1 failed (23,34) and required further optimisation to recover functional His::*Pf*PLSCR from inclusion bodies (Fig. 2A). The solubilised protein was refolded by rapid dilution prior to immobilised metal affinity chromatography and gel filtration. Optimal protein stability was achieved by using the zwitterionic surfactant Fos-Choline-12 as determined in a comprehensive detergent screen.

**Figure 2.**
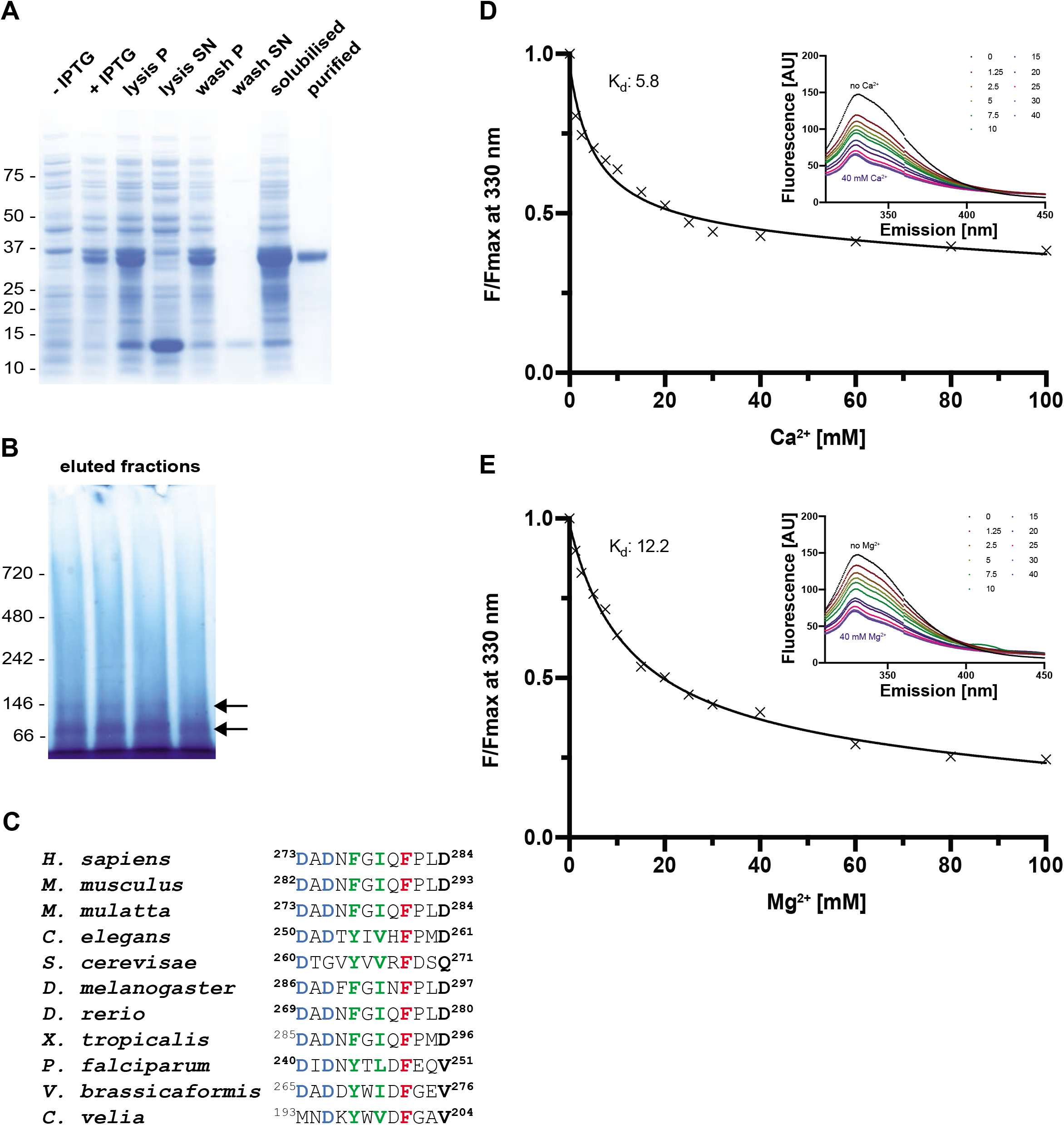
*Pf*PLSCR undergoes conformational changes in the presence of metal ions. A) SDS-PAGE of the expression and solubilisation of His::*Pf*PLSCR. The purified recombinant protein runs at the predicted molecular weight of ~ 37 kDa. Pellet and supernatant fractions are labelled as ‘P’ and ‘SN’, respectively. B) BN-PAGE of eluted fractions from the main peak reveals two protein species between the 66 kDa and 146 kDa protein markers (black arrows), indicating a possible dimerisation/oligomerisation of His::*Pf*PLSCR. C) Alignment of the putative Ca2+ binding regions from different PLSCR1 orthologues. Residues in positions 1, 3, 5, 7, 9 and 12 of the EF-hand-like motif in hPLSCR1 have been proposed to octahedrally coordinate the calcium ion (71) and are shown in bold. Highly conserved residues are highlighted in blue, hydrophobic residues (positions 5 and 7) in green and the strictly conserved phenylalanine residues (position 9) are shown red. D & E) Representative binding curves and calculated affinities of recombinant *Pf*PLSCR for Ca2+ and Mg2+ ions, respectively. The concentration-dependent decrease of fluorescence at λ 330 nm is plotted as F/Fmax with F= fluorescence measured in the presence of cations and Fmax= fluorescence under EGTA conditions. A selection of emission spectra is shown in the upper right corner.

A recent study provided first evidence for Ca^2+^-induced oligomerisation of hPLSCR1 which in turn activates PL scrambling (24). Hence, all steps of inclusion body solubilisation, protein refolding and purification of His::*Pf*PLSCR were carried out in the presence of 0.5 mM Ca^2+^ and Fos-Choline-12. Association of detergent micelles with membrane proteins is known to obscure the elution behavior in size-exclusion chromatography. We therefore performed BN-PAGE to assess the native (and potentially oligomeric) state of His::*Pf*PLSCR, which revealed two protein species suggestive of a dimeric and tetrameric form, respectively (Fig. 2B). Further attempts to determine the exact molecular mass of the protein species by size-exclusion chromatography multi-angle light scattering

(SEC-MALS) delivered ambiguous results (data not shown), most likely due to the presence of different oligomeric species in the protein solution as implicated by the BN-PAGE of individual fractions.

### *Pf*PLSCR is responsive to metal ions

The hPLSCR1 protein contains a single EF-hand like Ca^2+^ binding loop, which has been shown to bind Ca^2+^ and other divalent cations in a co-ordinated manner, thus inducing conformational changes of the protein (24,25,35). The unconventional Ca^2+^ binding motif of hPLSCR1 appears to be conserved in the parasite ortholog, including the residues in positions 1 (Asp^240^), 3 (Asp^242^) and 9 (Phe^248^) (Fig. 2C), all of which have been found to be crucial to the PL translocase activity of hPLSCR1 (20). A hydrophobic residue in the last position of the Ca^2+^ binding motif in *Plasmodium* spp. and Chromerida PLSCR orthologs replaces the aspartic acid residue in hPLSCR1 (and other orthologous proteins) (Fig. 2C). The presence of three tryptophan residues in *Pf*PLSCR enabled the use of intrinsic fluorescence measurements to analyse potential protein conformational changes of *Pf*PLSCR upon binding to Ca^2+^ and Mg^2+^ ions, respectively. Recombinant His::*Pf*PLSCR was therefore titrated with increasing amounts of the respective metal ion until saturation was achieved. Equilibrium dissociation constants were determined by using nonlinear regression analysis as described in (24).

His::*Pf*PLSCR consistently displayed an emission maximum at λ 330 nm. Addition of Ca^2+^ or Mg^2+^ ions resulted in a concentration-dependent decrease of fluorescence intensity with no shift in the emission peak of the protein (Fig. 2D-E). Similar observations were made for hPLSCR1 and hPLSCR2 exhibiting an emission maximum at λ 345 nm (24). The affinity of the recombinant parasite protein to the two metal ions tested was similar with dissociation constants (Kd) of 6.7 ± 0.6 mM (mean of n= 6 ± SD) for Ca^2+^ and 11.1 ± 1.7 mM (mean of n= 3 ± SD) for Mg^2+^ ions, respectively (Fig. 2D-E). *Pf*PLSCR therefore appears to have an approximately 4-fold higher affinity for Ca^2+^ and a 13-fold higher affinity for Mg^2+^ compared to the values determined for hPLSCR1 (24). Overall, our observations suggest a functional preservation of the Ca^2+^ binding site in *Pf*PLSCR and imply conformational changes of the protein in response to Ca^2+^ and Mg^2+^, albeit with low affinity as determined by binding constants in the mM range.

### *Pf*PLSCR mediates the transfer of PE and PS in reconstituted proteoliposomes

To investigate the putative PL translocating properties of *Pf*PLSCR, we reconstituted the recombinant protein into liposomes of different lipid compositions and performed ‘scramblase assays’ which allowed us to visualise and monitor PL translocase activity. Addition of sodium dithionate, which cannot cross the artificial membrane, to symmetrically labelled liposomes, reduces fluorescently labelled nitrobenzoxadiazole (NBD) lipids in the outer leaflet, resulting in an approx. 50% decrease of the initial fluorescence measured. Scrambling activity can be investigated by the ability of the putative *Pf*PLSCR to transfer NBD lipids from the inner to the outer leaflet by which their exposure to the reducing environment will lead to a further decrease in fluorescence (Fig. 3A).

**Figure 3.**
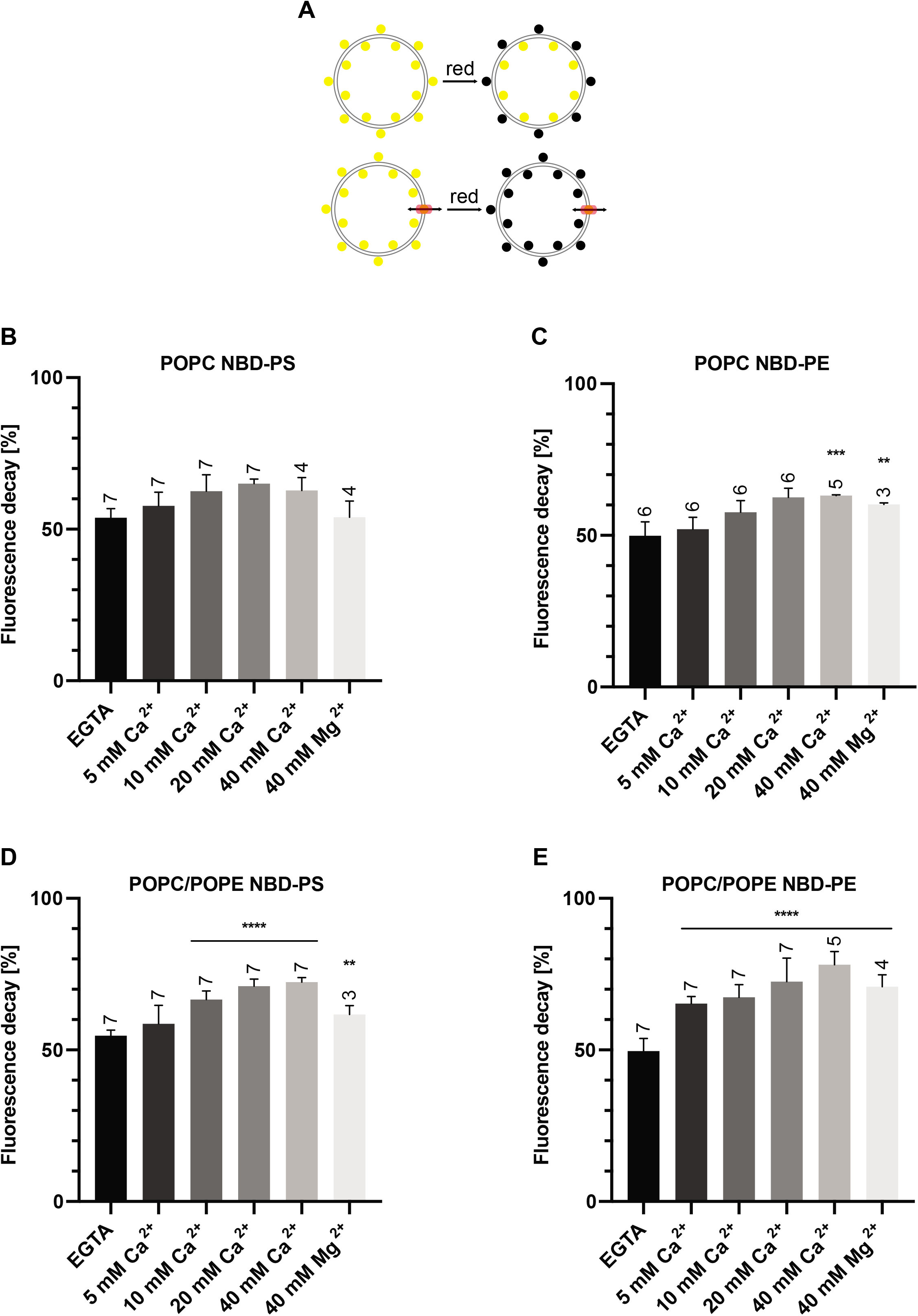
PL scramblase activity of recombinant *Pf*PLSCR in symmetrically labeled vesicles. **A)** Schematic of the ‘scramblase’ assay. **B & D)** Transbilayer movement of NBD-PS and **C & E)** NBD-PE in proteoliposomes of different lipid compositions. Scramblase activity (mean ± SD) is presented as the percentage of fluorescence decrease at 500 sec. Number of independent experiments is indicated above the error bars. Statistical significance of differences between activities measured in the presence of EGTA and divalent cations was calculated using unpaired t-tests (****, *p*<0.0001; ***, *p*=0.0001; **, *p*≤0.005).

No conclusive PL translocase activity of recombinant His::*Pf*PLSCR could be measured in proteoliposomes composed of PC only and NBD-PS (Fig. 3B & Fig. S2A). Some activity, however, was observed if NBD-PE was used as a substrate, leading to a further ~15% reduction in fluorescence and maximal catalytic function in the presence of 40 mM Ca^2+^ (Fig. 3C & Fig. S2B). No PL scrambling could be detected under EGTA conditions, presumably rendering the protein inactive as a consequence of ineffective oligomerisation. Translocase activity could also be observed in the presence of 40 mM Mg^2+^ but with reduced efficiency and comparable to the fluorescence decrease at 10 mM Ca^2+^ (Fig. 3C & Fig. S2B). Respective liposomes without *Pf*PLSCR displayed no further decrease in fluorescence beyond the 50% margin as expected (Fig. S2A-B).

In attempts to mimic the PL composition of the parasite’s membranes, an overall greater catalytic activity was noted in reconstituted proteoliposomes composed of PC and PE (70:30%), with a further decrease in fluorescence of maximally ~20% for vesicles labelled with NBD-PS (Fig. 3D & Fig. S2C) and ~30% for NBD-PE containing proteoliposomes (Fig. 3E & Fig. S2D). Phospholipid scrambling activity was enhanced by the increase of metal ion concentrations, reaching maximal activity at 20-40 mM Ca^2+^. Mg^2+^ activated PL scrambling could also be observed in these vesicles but once again with lower efficiency.

These data strongly suggest a metal ion dependent PL translocase function for *Pf*PLSCR with a preference for calcium ions and NBD-PE as a substrate as indicated by maximal activity under these conditions. A recent lipidomic analysis of the total lipid composition in asexual blood stage parasites showed that PC and PE accounted for 50% of the total lipid content (3). Inclusion of PE in reconstituted proteoliposomes resulted in an overall increase of His::*Pf*PLSCR mediated PL translocase efficiency, likely due to a more physiological environment allowing for optimal protein function.

### Generation of an inducible *Pf*PLSCR knock out line

Having demonstrated PL translocase activity for *Pf*PLSCR, we undertook a genetic approach to characterise the protein in *P. falciparum* parasites. We used a previously published genetic system (36) that modifies the endogenous *Pfplscr* locus by single-crossover homologous recombination using the selection-linked integration (SLI) strategy (37) and generated a conditional knock out (cKO) parasite line expressing C-terminally triple-HA tagged full length *Pf*PLSCR (hereafter referred to as *Pf*PLSCR-HA) under its endogenous promoter. Two synthetic loxPint modules (38) were inserted into the targeting plasmid, allowing for DiCre-mediated excision of the sequence encoding the C-terminal region Lys^217^-Lys^275^ of the protein in the optimised DiCre recombinase expressing B11 parasite line (39) (Fig. 4A). Transgenic parasites were retrieved after 2 weeks of initial WR99210 selection and a clonal population of integrant parasites was obtained 7 days after neomycin selection using SLI (37). Successful modification of the *Pfplscr* locus was confirmed by diagnostic PCR (Fig. 4B) and Western blot analysis giving rise to a ~ 37 kDa protein (doublet) band as predicted for endogenously tagged *Pf*PLSCR-HA (Fig. 4C). The growth rate of *Pf*PLSCR-HA expressing parasites was similar to that of the parental B11 line (Fig. 4D), suggesting no negative side effects of the modification to parasite viability.

**Figure 4.**
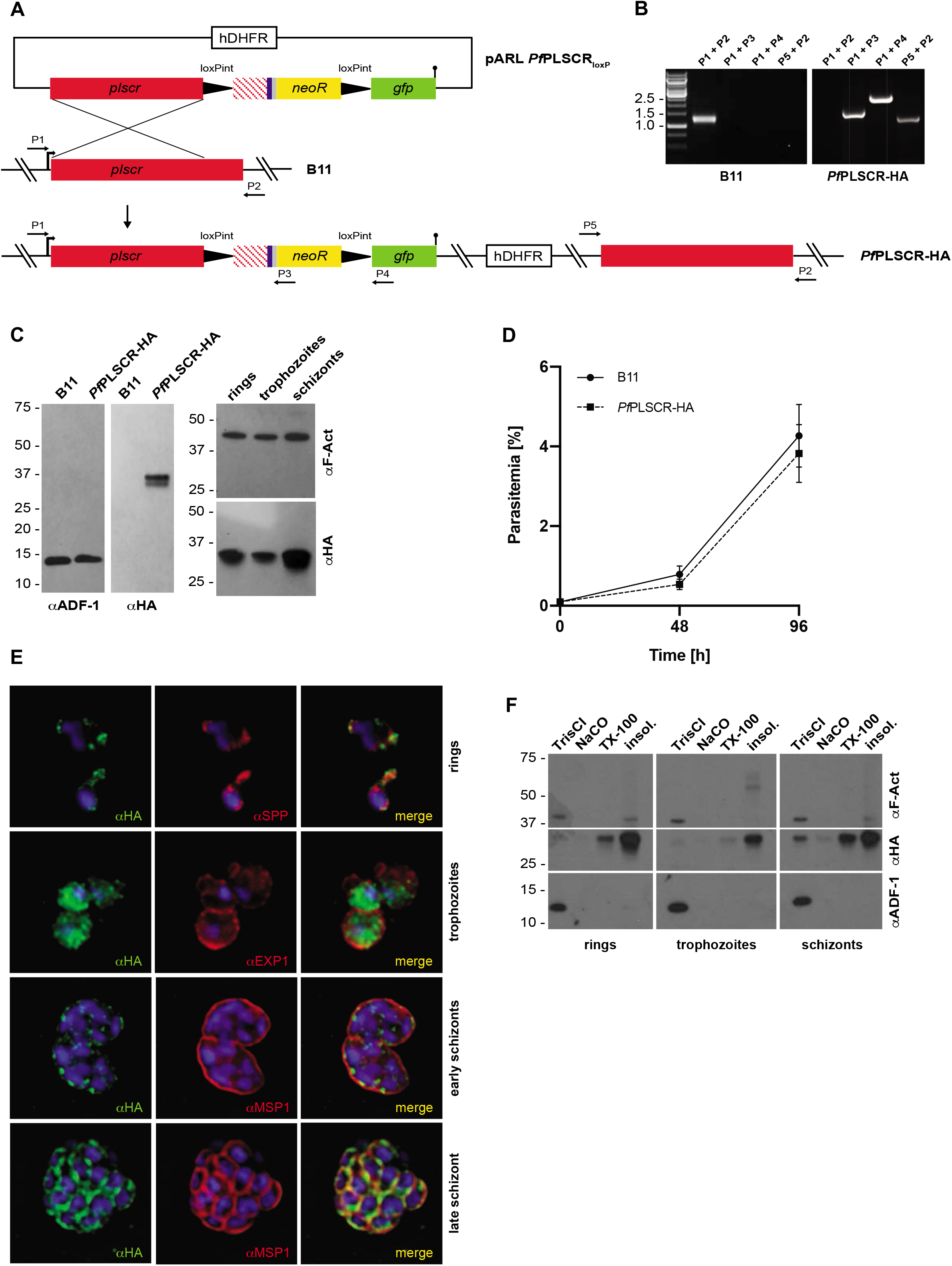
*Pf*PLSCR-HA is expressed throughout the asexual life cycle and localizes to membranous structures in the parasite. **A)** Schematic of the strategy to generate the conditional KO line *Pf*PLSCR-HA via SLI mediated single-crossover homologous recombination. The targeting plasmid contains the *Pfplscr* homology region (red), followed by two synthetic loxPint modules flanking the recodonised *Pfplscr* sequence (red stripes) with a triple-HA tag (blue), T2A skip peptide (gray) and neomycin resistance cassette (yellow). The GFP coding sequence (green) is followed by a stop codon (black bar). Numbered arrows indicate primers used for genotyping. **B)** Diagnostic PCR confirms successful integration into the *Pfplscr* locus by amplification of three distinctive fragments at the expected size and loss of the wild-type band in the *Pf*PLSCR-HA parasite line. **C)** Western blot analysis gives rise to a ~ 37 kDa protein band as predicted for *Pf*PLSCR-HA. The endogenously tagged protein is expressed throughout the intra-erythrocytic life cycle as indicated by the different parasite stages (hours post invasion): rings (4 ± 2 h), trophozoites (20 ± 4 h) and schizonts (44 ± 4 h). Anti-ADF (actin depolymerising factor) and anti-F-Actin antibodies are used as loading controls. **D)** Growth curves of parental B11 and *Pf*PLSCR-HA expressing parasites over two cycles. Parasitemias (as quantified by flow cytometry) were averaged from at least three biological replicates using blood from different donors and are presented as mean ± SD. **E)** Immunofluorescence analysis of *Pf*PLSCR-HA expressing parasites. The upper panel depicts two ring-stage parasites and co-labelling with anti-SPP (signal peptide peptidase) antibodies as an ER marker. Anti-EXP1 (exported protein 1) delineates the PVM surrounding three trophozoites in a multiply infected red blood cell. Lower two panels show deconvolved co-labelling with antibodies against MSP1 (merozoite surface protein 1) in late blood stage parasites. *Pf*PLSCR-HA co-localises with MSP1 at the plasma membrane in late schizonts. Parasite nuclei are stained with DAPI (blue). **F)** *Pf*PLSCR-HA remains strongly membrane-associated throughout the intra-erythrocytic life cycle as suggested by sequential extraction of the protein and its presence in the TX-100 insoluble fraction. The blot was cut and probed with anti-ADF as a control for soluble ADF protein and anti-F-Actin as a control for partially membrane-bound parasite actin.

### *Pf*PLSCR-HA localises to different membranous compartments throughout asexual development

The *Pfplscr* gene is transcribed throughout the intra-erythrocytic life cycle but is maximally expressed in late schizonts stages (40-48 h post invasion) (www.PlasmoDB.org). Immunoblot analysis of parasite protein lysates prepared at different timepoints post invasion confirmed the expression of *Pf*PLSCR-HA across the asexual life cycle (Fig. 4C). Immunofluorescence analysis revealed the localisation of the epitope-tagged protein to the periphery of individual merozoites in late schizonts as suggested by co-localisation with the merozoite surface protein MSP1 (Fig. 4E). The protein also localised to a peri-nuclear region in the early ring stages as evident by partial colocalisation with the ER-resident protein signal peptide peptidase (40), and redistributed to the cytoplasm and vesicular foci in early trophozoite and schizont stages (Fig. 4E). Several attempts to verify the localisation of *Pf*PLSCR-HA at the plasma membrane and other membranous compartments by immuno-EM proved inconclusive, likely due to the inaccessibility of the HA epitope being buried in the membrane. A solubility analysis, however, confirmed a tight membrane association of *Pf*PLSCR-HA throughout the asexual life cycle (Fig. 4F) supporting a possible PL translocase function in the membrane bilayer.

### *Pf*PLSCR is not essential for erythrocyte invasion and asexual development

Investigations into the nature of the C-terminal transmembrane helix of hPLSCR1 confirmed its function in inserting the protein into the bilayer of reconstituted liposomes and revealed an additional role in coordinating the metal ion, thus influencing scramblase activity (21,41). Two cholesterol-binding sites, composed of Ala^291^, Phe^298^, Leu^299^ and Leu^299^, Phe^302^, Glu^306^, respectively have also been identified and proposed to stabilise the C-terminal transmembrane helix in the phospholipid bilayer. The unusual Asp^301^ residue has further been suggested to play a role in scramblase activity (42).

To explore the specific role of *Pf*PLSCR in parasite development, rapamycin induced, DiCre-mediated excision of the ‘floxed’ sequence in the *Pf*PLSCR-HA line was anticipated to generate a mutant scramblase protein lacking the putative NLS, Ca^2+^-binding motif and C-terminal transmembrane helix (Lys^217^-Lys^275^ which includes the conserved Asp^268^ residue) and therefore PL translocase activity, which we hypothesised to be important for host cell invasion and/or intra-erythrocytic development (Fig. 5A). Diagnostic PCRs confirmed successful excision of the locus compared to DMSO-treated *Pf*PLSCR-HA parasites (Fig. 5B). Excision also removed the *neo-R* gene, enabling expression of the C-terminally truncated scramblase protein in fusion with GFP as evidenced by Western blot analysis (Fig. 5C-D). Live-cell fluorescence of the remaining GFP fusion protein was weak and not clearly distinguishable from background fluorescence but as predicted, plasma membrane localisation in merozoites was abolished as visualised by immuno-fluorescence analysis using anti-GFP antibodies in the conditional KO parasites (Fig. 5D). However, no significant growth or invasion defect could be detected for parasites expressing truncated *Pf*PLSCR in the induced KO compared to full length *Pf*PLSCR-HA in mock-treated parasites over a total period of 3 cycles as analysed by flow cytometry (Fig. 5E) or light microscopy of Giemsa-stained blood smears (data not shown). These findings suggest *Pf*PLSCR is dispensable for erythrocyte invasion and asexual parasite development.

**Figure 5.**
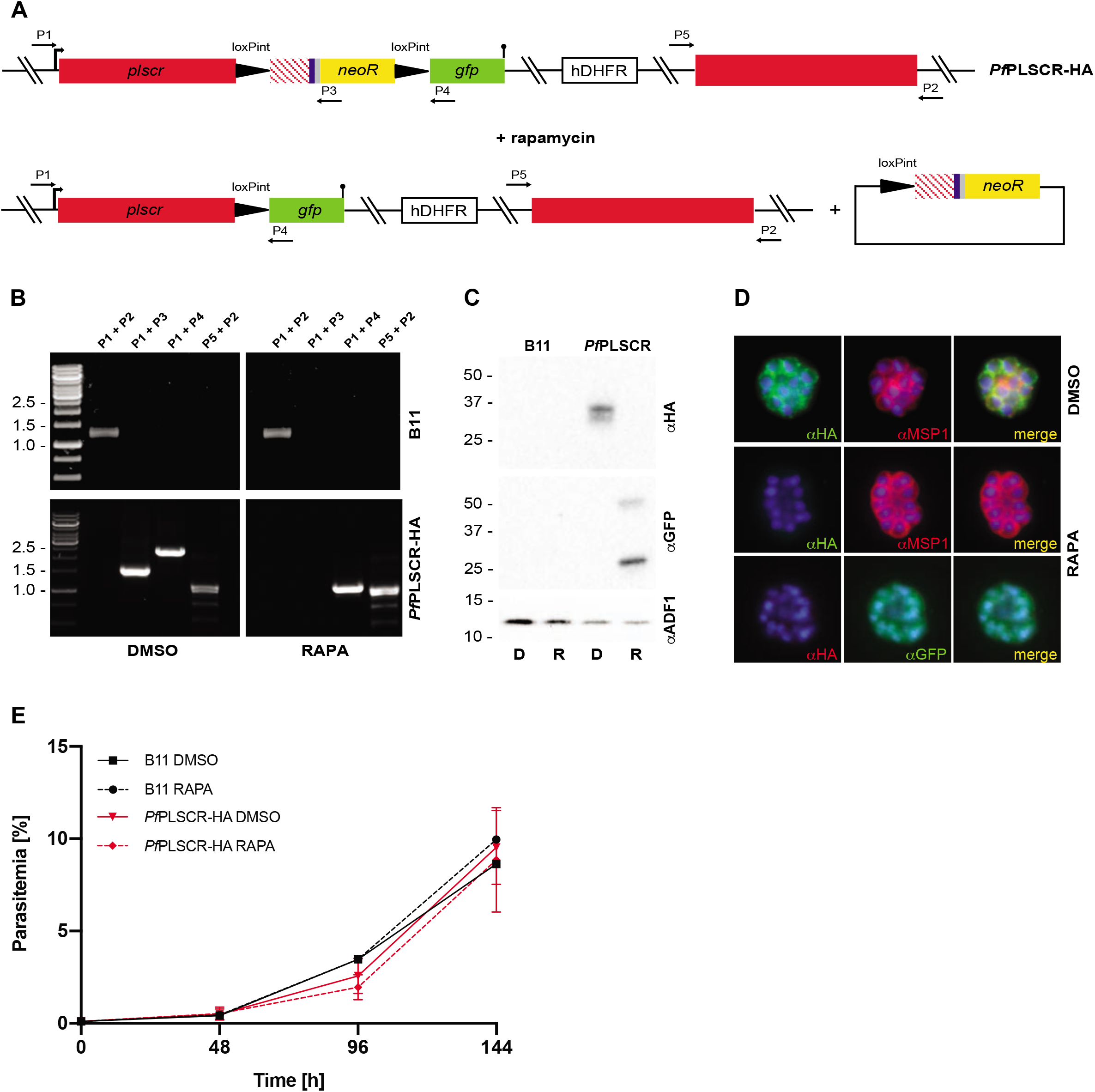
PL scrambling is not essential for red blood cell invasion and intra-erythrocytic parasite development. **A)** Schematic of rapamycin induced and DiCre-mediated excision of the floxed sequence in *Pf*PLSCR-HA. **B)** Diagnostic PCR confirms successful excision of the locus compared to DMSO-treated *Pf*PLSCR-HA parasites. **C)** The C-terminally truncated *Pf*PLSCR protein is expressed as a GFP fusion protein at the expected molecular weight of ~ 53 kDa in rapamycin-treated parasites. The ~ 27 kDa band is characteristic for GFP only. Anti-ADF antibodies are used as a loading control. **D)** Plasma membrane association is lost in parasites expressing the truncated *Pf*PLSCR-GFP fusion protein, which distributes throughout the parasite cytosol. **E)** Growth curves of rapamycin and DMSO-treated parasites over three cycles. Parasitemias were averaged from at least three biological replicates using blood from different donors and are presented as mean ± SD.

To exclude any residual activity of the truncated *Pf*PLSCR protein, we attempted a full KO of the *Pfplscr* locus using CRISPR-Cas9 by replacing the gene with a super folder GFP (sfGFP) expression cassette (Fig. 6A). Different combinations of the plasmid harbouring the Cas9 expression cassette (43), as well as one of three different guide RNAs tested, were co-transfected with the donor plasmid. One plasmid combination, containing guides 952497 and 952530, both in close proximity to the 5’ targeting flank, resulted in successful parasite recovery after 5 weeks post transfection as evident by sfGFP positive parasites (Fig. 6D). Replacement and loss of the *Pfplscr* gene with the sfGFP expression cassette was further confirmed by diagnostic PCR (Fig. 6B). A growth analysis of 3D7 wild-type versus *Pf*PLSCR KO parasites revealed no significant differences (Fig. 6C), further corroborating the redundancy of the PL scramblase protein in *Plasmodium* spp. asexual development.

**Figure 6.**
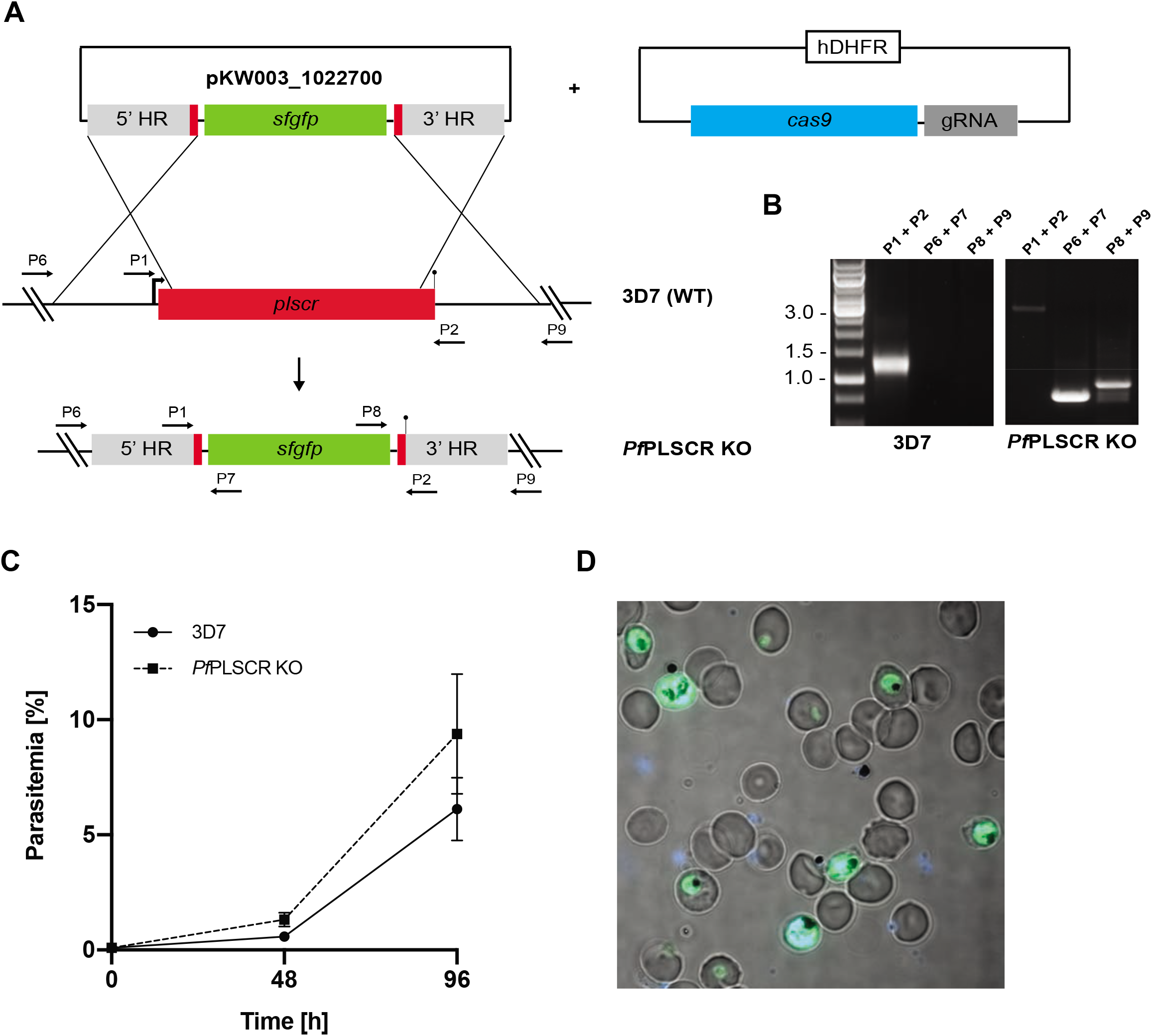
A full gene KO in 3D7 parasites supports the redundancy of *Pf*PLSCR in asexual parasite biology. **A)** Schematic of the CRISPR-Cas9 mediated KO strategy. Expression of sfGFP is driven under a constitutive BiP promoter. Numbered arrows indicate primers used for genotyping. **B)** Diagnostic PCR confirms successful replacement of the *Pfplscr* locus by the sfGFP expression cassette. **C)** Growth curves of wild-type 3D7 and *Pf*PLSCR KO parasites over two cycles. Parasitemias were averaged from at least three biological replicates using blood from different donors and are presented as mean ± SD. **D)** Live-cell image of *Pf*PLSCR KO parasites expressing sfGFP throughout asexual parasite development. Parasite nuclei are stained with DAPI (blue).

### *Pf*PLSCR is expressed in gametocytes

Transcriptomic data suggests that *Pf*PLSCR is also transcribed in the sexual stages of *P. falciparum* (www.PlasmoDB.org). A recent report on the role of a female gametocyte-specific ATP-binding cassette transporter (gABCG2) in regulating the metabolism of neutral lipids (44) prompted us to investigate the expression of *Pf*PLSCR-HA in gametocytes. Immunofluorescence analysis of stage III-IV gametocytes revealed different localisation patterns of the epitope-tagged protein. Localisations ranged from an intracellular distribution with more or less pronounced vesicular foci (Fig. 7A), to a peripheral localisation as indicated by co-labelling with Pfs16, a gametocyte specific protein of the PVM (Fig. 7A, last panel) and GAP45 (glideosome associated protein, Fig. 7B), a component of the IMC (inner membrane complex), which has also been identified in gametocyte stages (45). As the PVM, parasite plasma membrane and IMC cannot be distinguished at this resolution, a more detailed investigation at the ultrastructural level will be necessary to determine the exact location of *Pf*PLSCR-HA at the parasite’s periphery in gametocytes. However, imaging and immunoblot analysis confirm the expression of *Pf*PLSCR-HA in gametocytes (Fig. 7C) and, though negligible for asexual development, suggests that a possible role of the PL translocase in the sexual development of *Plasmodium* spp. parasites may yet be forthcoming.

**Figure 7.**
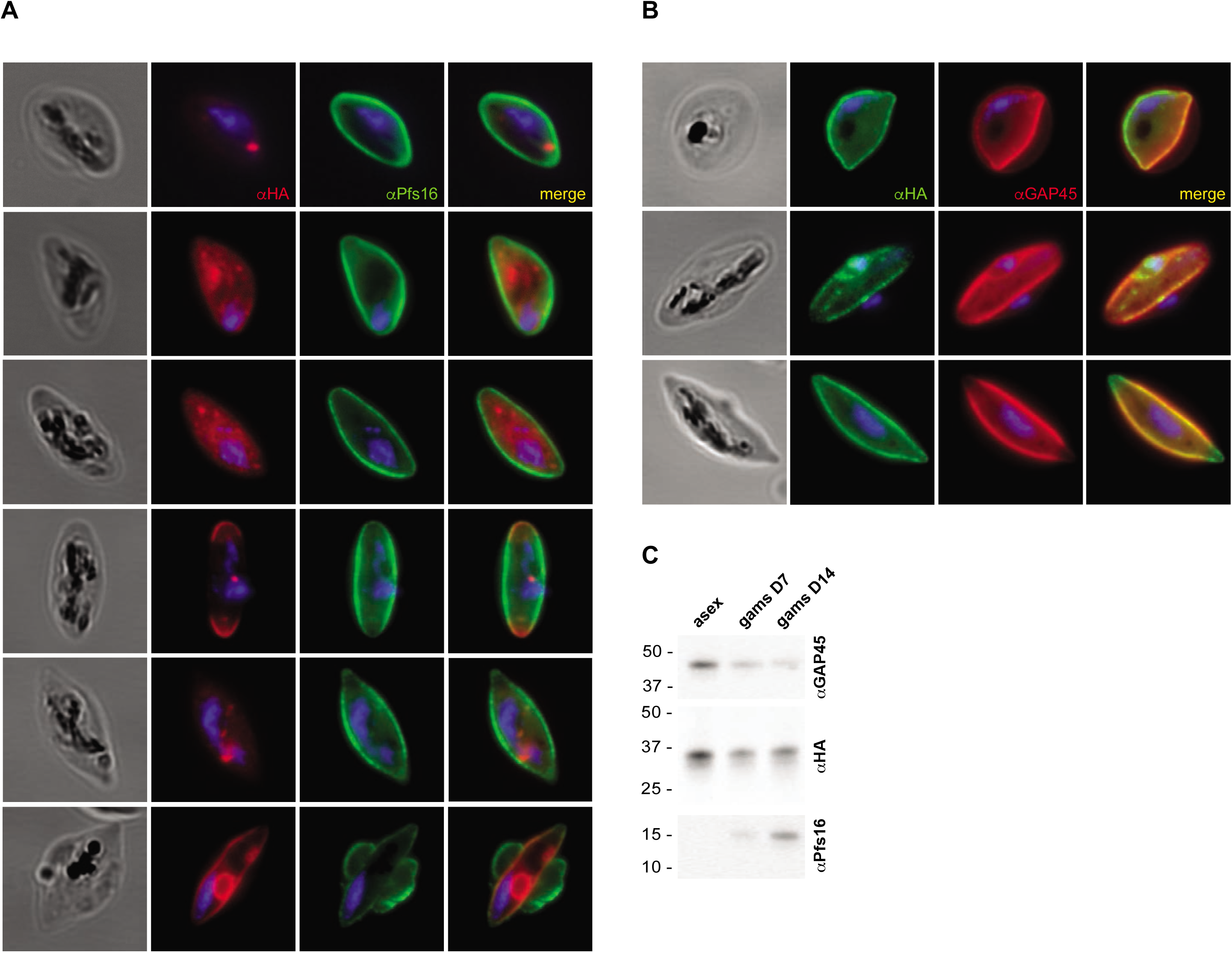
*Pf*PLSCR is expressed in gametocytes. **A)** Immunofluorescence analysis of stage III-IV gametocytes reveals different localisation patterns of the triple-HA tagged protein. Anti-Pfs16 is used as gametocyte specific antibody marking the PVM. Parasite nuclei are stained with DAPI. **B)** *Pf*PLSCR-HA co-localises with the IMC marker GAP45, which delineates the parasite periphery. **C)** Western blot analysis confirms *Pf*PLSCR-HA expression in gametocytes.

## Discussion

In this study, we identified and characterised a putative PL scramblase in the malaria parasite *P. falciparum*, which we hypothesised to play a role in erythrocyte invasion and/or asexual parasite development. A sequence comparison with the orthologous protein hPLSCR1 revealed a shorter proline-rich N-terminus in *Pf*PLSCR but the functional domains of the C-terminal portion appear to be conserved (Fig. 1B). Conservation in the C terminus is consistent with observations made for the four hPLSCR homologs and various other PLSCR orthologs found throughout the animal kingdom [reviewed in (30)]. The proline-rich domain (PRD) has been shown to mediate the subcellular localisation, Ca^2+^-dependent aggregation and therefore PL scrambling activity of hPLSCR1. The absence of this domain in hPLSCR2 renders the protein functionally inactive as a PL translocase but can be restored by addition of the hPLSCR1 PRD to the N-terminus of hPLSCR2 (24). The PXXP and PPXY motifs found in hPLSCR1, 3 and 4 are further thought to serve as potential binding sites for proteins containing SH3 and WW domains (14), scaffolds which recognize proline-rich regions in a variety of signaling and structural proteins.

Despite a shorter PRD and only one potential PXXP motif, intrinsic tryptophan fluorescence measurements on recombinant *Pf*PLSCR reveal Ca^2+^ and Mg^2+^ induced conformational changes, suggesting a functional preservation of the Ca^2+^ binding site. Residues in positions 1, 3, 5, 7, 9 and 12 of the EF-hand-like motif in hPLSCR1 have been proposed to octahedrally coordinate the Ca^2+^ ion (20) and are functionally conserved in the Ca^2+^ binding region of the parasite ortholog, with the exception of the terminal Val residue (Fig. 2C). Calculated dissociation constants in the mM range (Fig. 2D-E) indicate a low affinity for the divalent cations tested but are in accordance with the established values for hPLSCR1 and hPLSCR2 (24). BN-PAGE of His::*Pf*PLSCR purified in the presence of 0.5 mM Ca^2+^ suggests the formation of dimeric and/or oligomeric species of the protein (Fig. 2B), supporting the notion of Ca^2+^ mediated oligomerisation (24) and functionality of the reduced PRD region in *Pf*PLSCR. It should be noted that omission of Ca^2+^ in the purification of His::*Pf*PLSCR reduced aggregation during size-exclusion chromatography but rendered intrinsic fluorescent measurements inconclusive and the protein inactive in PL scramblase assays.

Maximal PL translocase activity of recombinant *Pf*PLSCR was measured in the presence of 40 mM Ca^2+^ and using NBD-PE as a substrate in proteo-liposomes composed of POPC and POPE (Fig. 3E & Fig. S2D), thus mimicking the lipidic environment of the parasite’s membranes (3). These experimental conditions resulted in an approx. 80% decrease of initial NBD-fluorescence recorded (50% quenching of fluorophores in the outer leaflet upon sodium dithionate treatment and a further 30% decrease due to translocase activity), which is well in line with established maximal activities of 80-85% in other proteins with PL scramblase function (46–48). The reported PL translocation rates for hPLSCR1 appear to be lower, most likely due to the use of NBD-PC as a substrate (with rates similar to those measured for NBD-PS in *Pf*PLSCR reconstituted vesicles) and/or insufficient reconstitution of recombinant hPLSCR1 into liposomes (20,23–25). Substrate preferences could also be observed in hPLSCR1 with higher catalytic activity for NBD-PE over NBD-PC (34) and in mitochondrial hPLSCR3, which preferentially translocates NBD-PE and NBD-PS, apart from cardiolipin (49). The exact mechanism of phospholipid translocation by the members of this PL scramblase family, however, remains to be resolved.

The generation of a conditional *Pf*PLSCR KO line enabled the investigation of triple-HA tagged full length *Pf*PLSCR in *P. falciparum* parasites. Expression of *Pf*PLSCR-HA under the endogenous promoter was observed throughout asexual parasite development with maximal expression in late schizont stages (Fig. 4C). Western blot analysis revealed a ~ 37 kDa HA-positive protein doublet, mainly present in late schizont stage protein samples, implying a possible modification of *Pf*PLSCR in the parasite. Human PLSCR1 was found to be phosphorylated by PKCδ in kinase assays (33) and differentially phosphorylated in Jurkat cells with a shift from serine phosphorylation in unstimulated to threonine phosphorylation in apoptotic cells (50). The authors further proposed an increased scramblase activity upon activation of PKCδ and phosphorylation of Thr^161^. However, no phospho-peptides could be identified for *Pf*PLSCR in a recent phospho-proteomic study of schizont samples (51), indicating a different nature of protein modification likely exists for *Pf*PLSCR.

*Pf*PLSCR-HA localises to different membranous compartments in asexual parasites (Fig. 4E-F). The redistribution of *Pf*PLSCR-HA from a perinuclear association in rings, to a more diffuse and vesicular dispersion in trophozoites and plasma membrane localisation in late schizonts resembles the observations made for hPLSCR1 which is trafficked to the plasma membrane of fibroblasts in the presence of palmitoylation but is found in the cytoplasm and nucleus of cells expressing mutant PLSCR lacking the five palmitoylation residues in the cysteine-rich motif ^184^CCCPCC^189^ (18). Mutation of the corresponding palmitoylation sites in hPLSCR3 and hPLSCR4 also results in nuclear redistribution (52,53). Nuclear import of hPLSCR1 and hPLSCR4 has been proposed to be mediated by a minimal non-classical nuclear leader sequence (NLS) and to occur via the α/β importin pathway (19,53,54). A subsequent study demonstrated transcriptional regulation of the IP3R1 receptor by hPLSCR1 (17), further supporting a potential nuclear function of hPLSCR1. *Pf*PLSCR possesses a predicted bi-partite NLS (Fig. 1B) and has been identified as a putative palmitoyl protein (55), concordant with the prediction of two palmitoylation sites at ^172^CC^173^ and ^194^CC^195^ [http://csspalm.biocuckoo.org (27)]. While this needs to be experimentally proven, stage-specific palmitoylation of *Pf*PLSCR could explain the differential localisation and presence of the discrete doublet for *Pf*PLSCR-HA.

The biochemical verification of PL translocase activity, maximal expression of *Pf*PLSCR-HA in late schizont stages, the localisation to the plasma membrane in merozoites and the unique conservation of this (otherwise ubiquitously present) protein in *Plasmodium* spp. but no other apicomplexan parasite reinforced our hypothesis of a role for *Pf*PLSCR in the infection of the host erythrocyte. Two different gene knockout approaches, however, revealed no invasion or intra-erythrocytic growth defect in the generated knockout lines (Fig. 5E & 6C). Plasma membrane association of *Pf*PLSCR is abolished in late stage parasites expressing the mutant *Pf*PLSCR GFP fusion protein in the conditional KO line (Fig. 5D) and supports an anchoring role of the predicted C-terminal transmembrane region in *Pf*PLSCR. Complete deletion of the gene in full *Pf*PLSCR KO parasites (Fig. 6) confirmed the redundancy of *Pf*PLSCR in asexual parasite development. A search of the PlasmoGEM database focused on the genetic manipulation of the rodent malaria parasite *P. berghei* also affirmed the rodent ortholog PBANKA_0506900 to be dispensable for intra-erythrocytic development, corroborating our results (https://plasmogem.sanger.ac.uk/).

While the biological function and preservation of *Pf*PLSCR remains elusive, expression of *Pf*PLSCR-HA in gametocytes (Fig. 7) may indicate a role in other stages of the parasite lifecycle or transitions between them. Another intriguing finding is the presence of orthologs in the closely related alveolates *Chromera velia* and *Vitrella brassicaformis*. Though the algae have not been found to sexually reproduce, they do form elongated zoospores with two heterodynamic flagella of unknown function (33), which resemble exflagellated male gametocytes of *P. falciparum* parasites. Recent reports on a female gametocyte-specific ATP-binding cassette transporter (gABCG2) regulating the numbers of gametocytes and accumulation of neutral lipids (44), changes in the lipid profile during sexual parasite development (3,56) and the involvement of host cell lyso-phosphatidylcholine in mediating sexual commit-ment and gametocytogenesis in *P. falciparum* parasites (57) and host phosphatidylcholine in mediating liver stage infection (58) highlight the emerging role of lipids and their regulators throughout the parasite’s life cycle and underpin the notion of a putative function for *Pf*PLSCR in the sexual stages of *Plasmodium* spp. parasites which encourages and gives substance to further exploration. Various cellular activities have been demonstrated and proposed for hPLSCR1, though the precise biological function of this multifaceted lipid effector and its counterpart in *Plasmodium* spp. remains to be unravelled.

## Materials and Methods

### Recombinant protein expression and inclusion body solubilisation

Codon-optimised full-length *Pfplscr* was synthesised as a gBlock^®^ (Integrated DNA Technologies) and cloned into the BamHI/XhoI site of the pPROEX HTa vector (Invitrogen), allowing the expression of recombinant *Pf*PLSCR with an N-terminal hexa-histidine affinity tag (His::*Pf*PLSCR) in *E. coli* BL21 (DE3) cells. Protein expression was induced with 1 mM IPTG for 14-16 h at 16°C. Bacterial cells were pelleted and lysed in lysis buffer [250 mM NaCl, 0.5 mM CaCl2, 1 mM MgCl2, 50 mM HEPES pH 7.5] using a high-pressure cell disruptor (Constant Systems). Insoluble material was then centrifuged at 30,000 g (30 min, 4°C) and washed twice with wash buffer [0.5 M NaCl, 0.5 mM CaCl2, 1 mM MgCl2, 50 mM HEPES pH 7.5] by sonication and centrifugation as described above. Inclusion bodies were solubilised in solubilisation buffer [250 mM NaCl, 0.5 mM CaCl2, 1 mM TCEP, 1 % SDS, 25 mM HEPES pH 7.5] under gentle agitation over night at 25°C and the soluble fraction collected by centrifugation at 30,000 g (30 min, 25°C). All buffers contained cOmplete™ EDTA-free protease inhibitors (Roche).

### Refolding and purification of His::*Pf*PLSCR

Recovered His::*Pf*PLSCR protein was refolded by rapid dilution into IMAC buffer A [250 mM NaCl, 0.5 mM CaCl2, 1 mM TCEP, 0.1% Fos-choline-12 (n-Dodecyl-phosphocholine, Anatrace), 25 mM HEPES pH 7.5], resulting in a final SDS concentration of 0.1%. Refolded protein was filtered through a 0.45 μm filter prior to immobilised metal affinity chromatography (IMAC) using a HisTRAP™ High Performance column (GE Healthcare), which was equilibrated with IMAC buffer A.

The loaded protein column was washed with several column volumes of wash buffer [IMAC buffer A + 0.4% SDS] and IMAC buffer A. The protein was eluted in a linear imidazole gradient using IMAC buffer B [IMAC buffer A + 1 M imidazole]. Fractions containing the desired protein were concentrated (Amicon Ultracel 100 K, Millipore) and further purified by size-exclusion chromatography (SEC) using a Superdex S200 10/300 GL column (GE Healthcare) in SEC buffer [250 mM NaCl, 0.5 mM CaCl2, 1 mM TCEP, 0.1% FC-12, 25 mM HEPES pH 7.5]. All buffers contained cOmplete™ EDTA-free protease inhibitors (Roche). Purity of the protein was assessed by SDS-PAGE using NuPAGE™ Novex^®^ Bis-Tris gels and blue native PAGE was performed by using the NativePAGE™ Novex^®^ Bis-Tris Gel System and G250 additive (Life Technologies).

### Intrinsic tryptophan fluorescence

Intrinsic tryptophan fluorescence of recombinant protein was measured using a Cary Eclipse Spectrophotometer (Agilent) and Jasco FP-8500 Spectrofluorometer, respectively. Cuvettes were washed with Helmanex™ III and rinsed with ion-free water before each use. Fresh stocks of CaCl2 and MgCl2 were prepared using ion-free water for metal ion titration and measurements in reconstitution buffer [100 mM NaCl, 10 mM HEPES, pH 7.5] (34) containing either 2 mM EGTA (for cation free conditions) or 2 mM EGTA and the desired metal ion concentration as calculated by using the Maxchelator programme [https://somapp.ucdmc.ucdavis.edu].

Five μg of purified recombinant protein were used per emission scan (in either duplicate or triplicate measurements per condition and experiment). Intrinsic fluorescence emission spectra were recorded from 310 to 470 nm at 25°C and an excitation wavelength of 295 nm. Emission and excitation bandwidths were set to 5 nm and 10 nm, respectively. Equilibrium dissociation constants were determined by using the nonlinear regression analysis tool and one-site total binding equation (GraphPad Prism) as described in (24).

### Reconstitution of recombinant His::*Pf*PLSCR into liposomes

The following lipids were obtained from Avanti Polar Lipids to prepare liposomes: 1-palmitoyl-2-oleoyl-sn-glycero-3-phosphocholine (POPC); 1-palmitoyl-2-oleoyl-sn-glycero-3-phosphoethanolamine (POPE); 1,2-di-oleoyl-sn-glycero-3-[(N-(5-amino-1-carboxypentyl) iminodiacetic acid) succinyl] nickel salt (18:1 DGS-NTA (Ni)); 1,2-dioleoyl-sn-glycero-3-phospho-L-serine-N-(7-nitro-2 - 1,3-benzoxadiazol-4-yl) (18:1 NBD-PS) and 1,2-dioleoyl-sn-glycero-3-phosphoethanolamine-N-(7-nitro-2-1,3-benzoxadiazol-4-yl) (18:1 NBD-PE).

To prepare liposomes, dried lipid films of either PC only or PC/PE (70:30% w/w), supplemented with 0.1% DGS-NTA (Ni) and either 0.25% NBD-PE (w/w) or 0.25% NBD-PS (w/w), respectively, were hydrated at 3 mg/ml total lipid in reconstitution buffer and incubated for 1 h at 37°C under agitation. The emulsion was then sonicated in a sonicator bath for 1 min and subjected to five freeze-thaw cycles. Uniformly sized ULVs were obtained using a lipid extruder mounted with a 100 nm membrane (Avanti Polar Lipids).

To prepare proteoliposomes, liposomes were destabilised with n-Dodecyl-β-D-Maltopyranoside (DDM, Anatrace) at 1:8 molar ratio (detergent to lipid) for 15 min at 25°C prior to the addition of purified His::*Pf*PLSCR (5 μg protein per mg of lipid). Aliquots of detergent treated liposomes were taken aside, matching volumes of SEC buffer were added instead of protein and liposome only controls were treated as proteoliposomes which were formed under gentle agitation and incubated at 25°C for 24 h. Detergent was removed by adding excess amounts (~ 100-fold) of reconstitution buffer and vesicles were collected by ultracentrifugation at 150,000 g (1 h, 25°C) and resuspended in reconstitution buffer containing 2 mM EGTA and at a lipid concentration of ~ 10 mg/ml.

### Scramblase assay

Scramblase activity was measured using a Cary Eclipse Spectrophotometer (Agilent) and Jasco FP-8500 Spectrofluorometer, respectively. Stirring cuvettes and bars were rinsed with Helmanex™ III and ion-free water before each use. Five μl of proteoliposome (or liposome only) suspension were diluted in 1 ml of reconstitution buffer containing either 2 mM EGTA or 2 mM EGTA and the desired metal ion concentration and stirred at 25°C. Emission spectra from 485 to 600 nm were recorded at 25°C and an excitation wavelength of 470 nm to determine the exact emission maxima of the NBD moieties (533 nm for NBD-PE and 546 nm for NBD-PS). Initial fluorescence was recorded for 100 sec prior to the addition of freshly prepared sodium dithionite (in unbuffered 0.5 M Tris) to a final concentration of 30 mM. Fluorescence decay was recorded for the next 400 sec and maximal activity was determined by dividing F= fluorescence at Time= 500 sec by Fmax= fluorescence prior to addition of 30 mM sodium dithionite. Traces were fitted using the ‘Plateau followed by one phase decay’ equation in GraphPad Prism.

### Transfection constructs

We used a recently published conditional gene deletion vector (36) to generate pARL *Pf*PLSCR-HA (Fig. 3A). A targeting fragment composed of the full length *Pfplscr* coding sequence followed by a triple hemagglutinin tag (HA), as well as two synthetic loxPint modules (38) flanking the recodonised sequence encoding the C-terminal protein region Lys^217^-Lys^275^ was synthesised (Geneart^®^) and cloned into pARL FIKK10.1 :loxPint: HA (a kind gift from the Treeck lab) by removing the *fikk10.1* gene via BglII/PstI.

Full *Pfplscr* gene disruption was attempted using CRISPR-Cas9 and by replacing the gene locus with a superfolder green fluorescent protein (sfGFP) expression cassette (Fig. 6A). To generate the donor plasmid pKW003_1022700, primers 1022700 5F and R were used to amplify the 572 bp 5’ homology flank and primer pair 1022700 3F and R was used to amplify the 673 bp 3’ homology flank (KOD Hot Start DNA Polymerase, Merck Millipore) which were cloned on either side of the sfGFP expression cassette in pkiwi003 (Ashdown et al., *in press*) using appropriate restriction sites (Table S1). The pDC2-cam-coCas9-U6.2-hDHFR vector (43) was used to clone in either of the guide RNAs 951969, 952497 and 952530, which were identified by using the web tool CHOPCHOPv2 [http://chopchop.cbu.uib.no (59)]. Therefore, two complimentary oligos (of each guide) with 5’ TATT (oligo 1) and 5’ AAAC (oligo 2) overhangs were annealed for ligation into the BbsI digested and dephosphorylated pDC2-cam-coCas9-U6.2-hDHFR vector (a kind gift from the Lee lab).

### Parasite culture and transfection

*P. falciparum* asexual parasites were cultured in RPMI 1640 medium containing 0.5% w/v AlbuMax II^®^ (Gibco) and at 4% haematocrit using human erythrocytes (blood group 0^+^) according to standard procedures (60). Ring-stage parasites were synchronised using sorbitol treatment. Tightly sychronised DiCre-expressing B11 parasites (39) were transfected with 100 μg of purified pARL *Pf*PLSCR-HA plasmid and initially selected with 2.5 nM WR99210 (Jacobus Pharmaceuticals) to obtain transgenic parasites harbouring the episomal plasmid. Integration into the genomic locus was achieved by G418 treatment (400 μg/ml) to select for integrants as described in (37).

DiCre-mediated recombination was induced by treating synchronised ring-stage parasites with 100 nM rapamycin (Sigma) in 1% (v/v) DMSO for 14-16 h to achieve complete excision. In parallel, parasites were mock-treated with 1% (v/v) DMSO as a negative control. Treated parasites were transferred into 96-well plates at 0.1% parasitemia (in triplicate) containing fresh red blood cells at 0.1% haematocrit and were allowed to proceed to the next ring-stage cycle for flow cytometry. Samples for gDNA extraction and PCR amplification, Western blot and immuno-fluorescence analysis were taken towards the end of the treatment cycle or subsequent cycles.

Ring-stage 3D7 parasites were co-transfected with different combinations of either guide RNA (and Cas9) containing plasmid 951969, 953497 and 953530, as well as with pKW003_1022700 donor plasmid (50 μg each). Drug selection with WR99210 was carried out for 6 days post transfection to ensure sufficient expression of the Cas9 nuclease. Rapid loss of the donor plasmid was assumed in the absence of any selectable marker and sfGFP positive parasites would therefore only be obtained upon successful replacement of the *Pfplscr* gene with the sfGFP expression cassette.

### *P. falciparum* gametocyte culture

Asexual parasite cultures were adapted to gametocyte culture media [RPMI 1640 medium containing 5% pooled human serum, 5% AlbuMax II^®^, 4% sodium bicarbonate and 3.5% Hypoxanthine-Thymidine supplement (Sigma-Aldrich)] prior to the induction of gametocytes. Gametocyte cultures were seeded from synchronised asexual cultures at high ring-stage parasitemia (~ 5-10%) and 4% haematocrit and were maintained at 37°C under 3% O2/5%°CO2/93% N2 gas (BOC, UK). Spent culture media was replaced daily for 14 days and cultures were treated with heparin (20 U/ml) on day 4 post induction for four days to kill uncommitted asexual parasites.

### Solubility analysis

Stage-specific infected RBCs were lysed with 0.02% saponin/PBS. Released parasite material was washed three times with PBS (containing protease inhibitors) prior to hypotonic lysis with 5 mM TrisCl (pH 8.0) for 1 h on ice. The soluble protein fraction was collected by ultracentrifugation at 100,000 g for 30 min at 4°C. The pellet was resuspended in 100 mM Na2CO3 (pH 11.2) to extract peripheral membrane proteins for 1 h on ice. The carbonate-soluble fraction was collected as above and the carbonate-insoluble pellet was needle-passed with 1% Triton X-100/PBS and incubated at 37°C for 30 min to extract integral membrane proteins. Triton X-100 soluble and insoluble fractions were separated as above and equal amounts of all fractions were analysed by immunoblotting.

### Western blot analysis

For immunoblots, infected RBCs were lysed and washed as above. Parasite material was resuspended in 1x SDS sample buffer. Extracted proteins were separated on NuPAGE™ Novex^®^ 4-12% Bis-Tris protein gels in MES buffer (Life Technologies) and transferred onto nitrocellulose membranes using the iBlot^®^ system (Life Technologies). Rabbit anti-HA (clone C29F4, Cell Signaling) was diluted 1:4,000; mouse anti-HA (clone 12CA5, Roche), mouse anti-GFP (clones 7.1 and 13.1, Roche), rabbit anti-ADF1 (61), rabbit anti-F-Actin (62) and mouse anti-Pfs16 (a kind gift from Professor Robert Sauerwein) were diluted 1:1,000 in 5% skim milk/PBS (w/vol). Horseradish peroxidase-conjugated goat anti-rabbit and goat anti-mouse antibodies were used as secondary antibodies (Jackson IR) and diluted 1:10,000 in 5% skim milk/PBS.

### Immunofluorescence analysis and imaging

Infected RBCs were fixed with 4% paraform-aldehyde and 0.0075% glutaraldehyde in PBS for 30 min, permeabilised with 0.1% Triton X-100/PBS for 10 min, and blocked in 3% BSA/PBS prior to immuno-labelling with the following primary antibodies diluted in 3% BSA/PBS: Rabbit anti-HA (1:1,000); mouse anti-HA (1:500), mouse anti-GFP (1:100), rabbit anti-MSP1 (1:500, (63)), rabbit anti-SPP (1:100, (40)), rabbit anti-EXP1 (1:100), mouse anti-Pfs16 (1:1000) and rabbit anti-GAP45 (1:500, (64)). Appropriate Alexa Fluor^®^ conjugated secondary antibodies (Life Technologies) and DAPI (4’,6-diamidino-2-phenylindole) were diluted 1:4,000 in 3% BSA/PBS. Images were captured using a Nikon Ti Microscope and OrcaFlash4.0 digital camera and were processed with Fiji (65). Z-stacks were deconvolved using the EpiDEMIC plugin with 80 iterations in Icy (66).

### Flow cytometry

Tightly synchronised ring-stage parasites were diluted to 0.1% parasitemia (in triplicate) for growth analysis and transferred into 96-well plates containing fresh red blood cells at 0.1% haematocrit. Parasites were allowed to proceed to the next ring-stage cycle and were incubated with SYBR Green (1:10,000, Sigma) for 10 min at 37°C in the dark. Stained cells were washed three times with PBS prior to collection by flow cytometry using a BD LSRFortessa™. A 100,000 cells were counted, FCS-A vs SSC-A was used to gate for red blood cells, FCS-A vs FCS-H for singlets and SYBR-A vs FCS-H was applied to gate for infected red blood cells (Fig. S2). Flow cytometry data was analysed in FlowJo. Biological replicates are indicated in the figures.

## Acknowledgments

We thank Matt Child for critically reading the manuscript, Moritz Treeck (Francis Crick Institute, UK) and Marcus Lee (Wellcome Sanger Institute, UK) for providing plasmids, as well as Robert Sauerwein (Radboud UMC), Brendan Crabb (Burnet Institute) and Alan Cowman (Walter & Eliza Hall Institute) for provision of antibodies. We are indebted to Simone Kunzelmann and the Structural Biology STP at the Francis Crick Institute for access to the Jasco FP-8500 Spectrofluorometer and to Marc Morgan at the Imperial College X-ray Crystallography Facility for help with SEC-MALS analysis and crystallisation trials. This work was supported by a Wellcome Investigator Award to JB (100993/Z/13/Z).

## Author contributions

S.H. conceived the project, designed, performed and coordinated the experiments as well as analysed and interpreted the data. M.C. optimised recombinant protein expression and purification, as well as D.M. who also provided guidance with the biochemical assays. D.C. assisted in recombinant protein expression and purification. S.J. assisted in preparing *P. falciparum* gametocyte cultures and samples. J.M.G. and J.B. provided scientific and advisory support. S.H. wrote the manuscript. All authors contributed to editing and approving the manuscript. The authors declare no conflicts of interest.

## Supplementary Information

**Figure S1.**
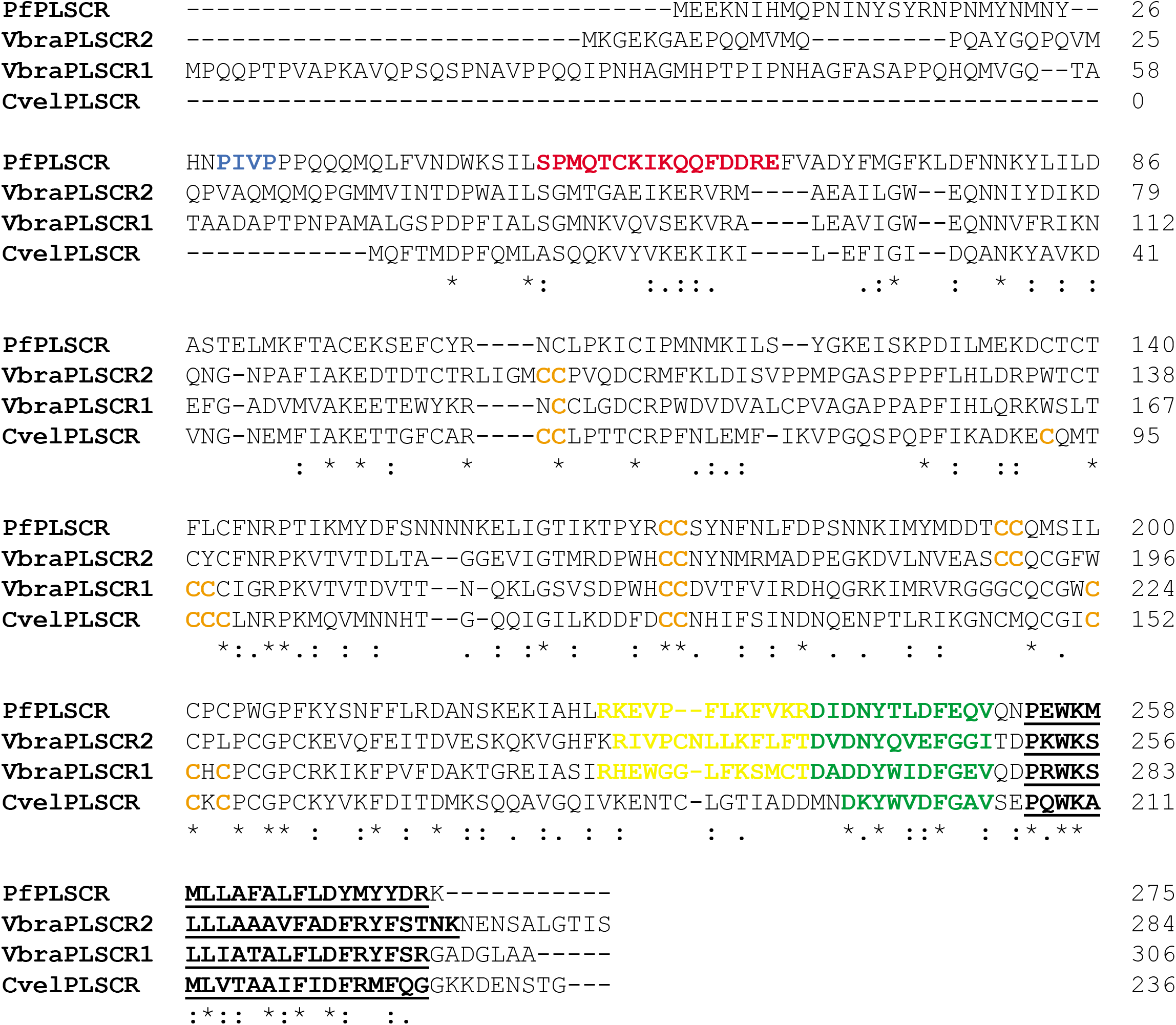
*Pf*PLSCR is conserved in unicellular algae. Alignment of *Pf*PLSCR with the orthologous proteins from the closely related *Chromera velia* and *Vitrella brassicaformis* algae. Similar residues are marked by colons (:) and identical residues are indicated by stars (*). Protein sequences were aligned with Clustal Omega (67). Predicted bipartite nuclear leader sequences are highlighted in yellow, putative Ca2+ binding regions in green and the C-terminal transmembrane helices are underlined. The cut-off score was set to 4.0 for the prediction of palmitoylation sites (68) highlighted in orange.

**Figure S2.**
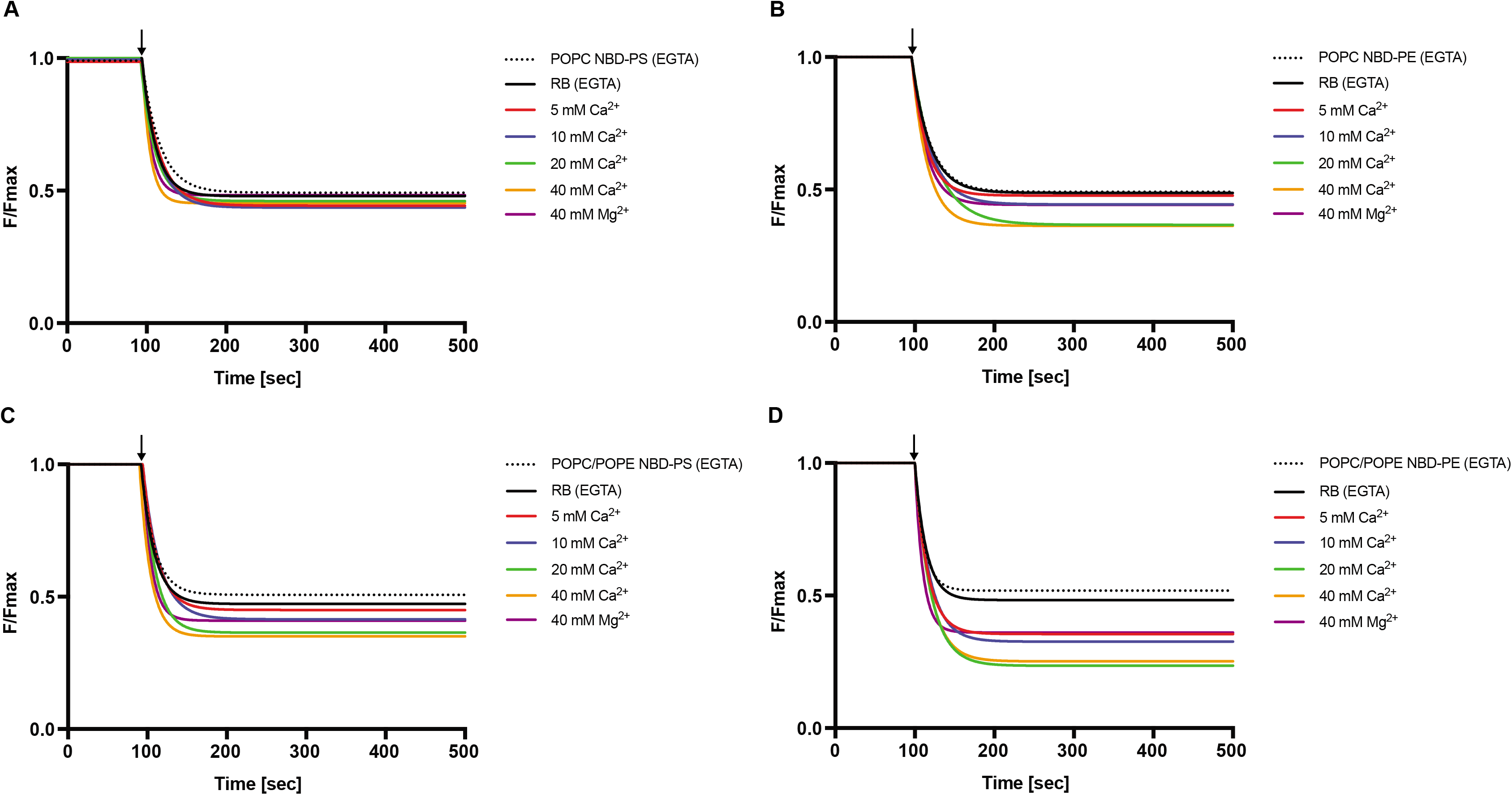
PL scrambling by recombinant *Pf*PLSCR. Representative traces are shown for each PL composition. The translocation of **A & C)** NBD-PS and **B & D)** NBD-PE in *Pf*PLSCR containing proteoliposomes was recorded in the presence and absence of Ca2+ and Mg2+ ions, respectively. Protein-free liposomes are denoted by dotted lines. The decay of fluorescence is plotted as F/Fmax with F= fluorescence at Time [sec] and Fmax= fluorescence prior to addition of 30 mM sodium dithionite (indicated by arrows at 100 sec).

**Figure S3.**
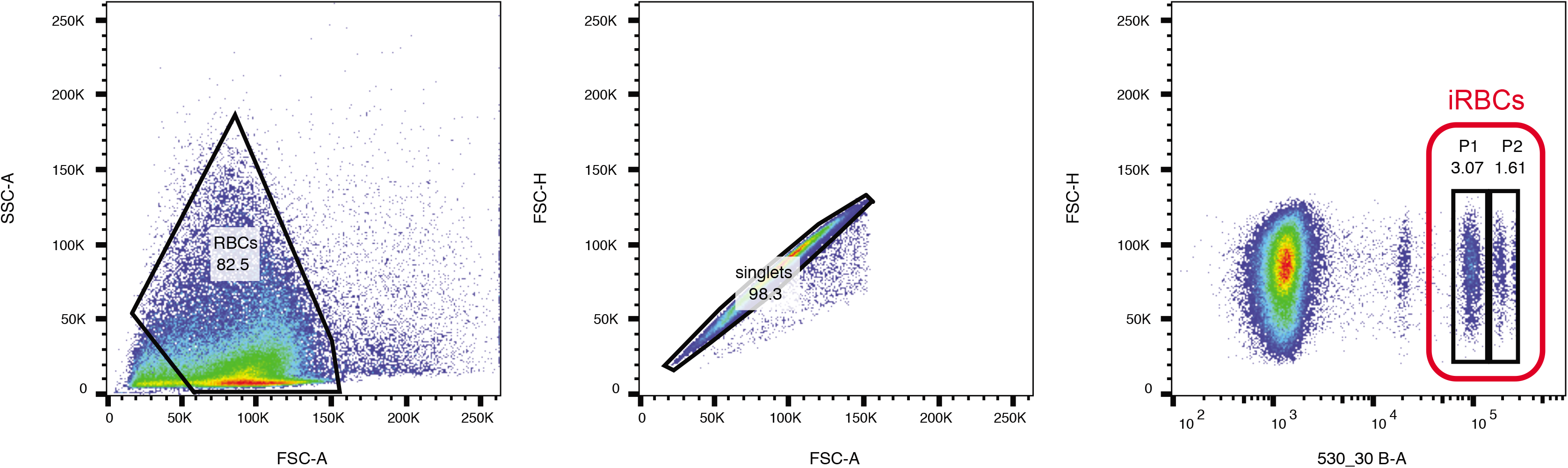
Gating strategy used to analyse flow cytometry data. The populations P1 and P2 denote single and multiply ring-stage infected red blood cells.

**Table S1.**
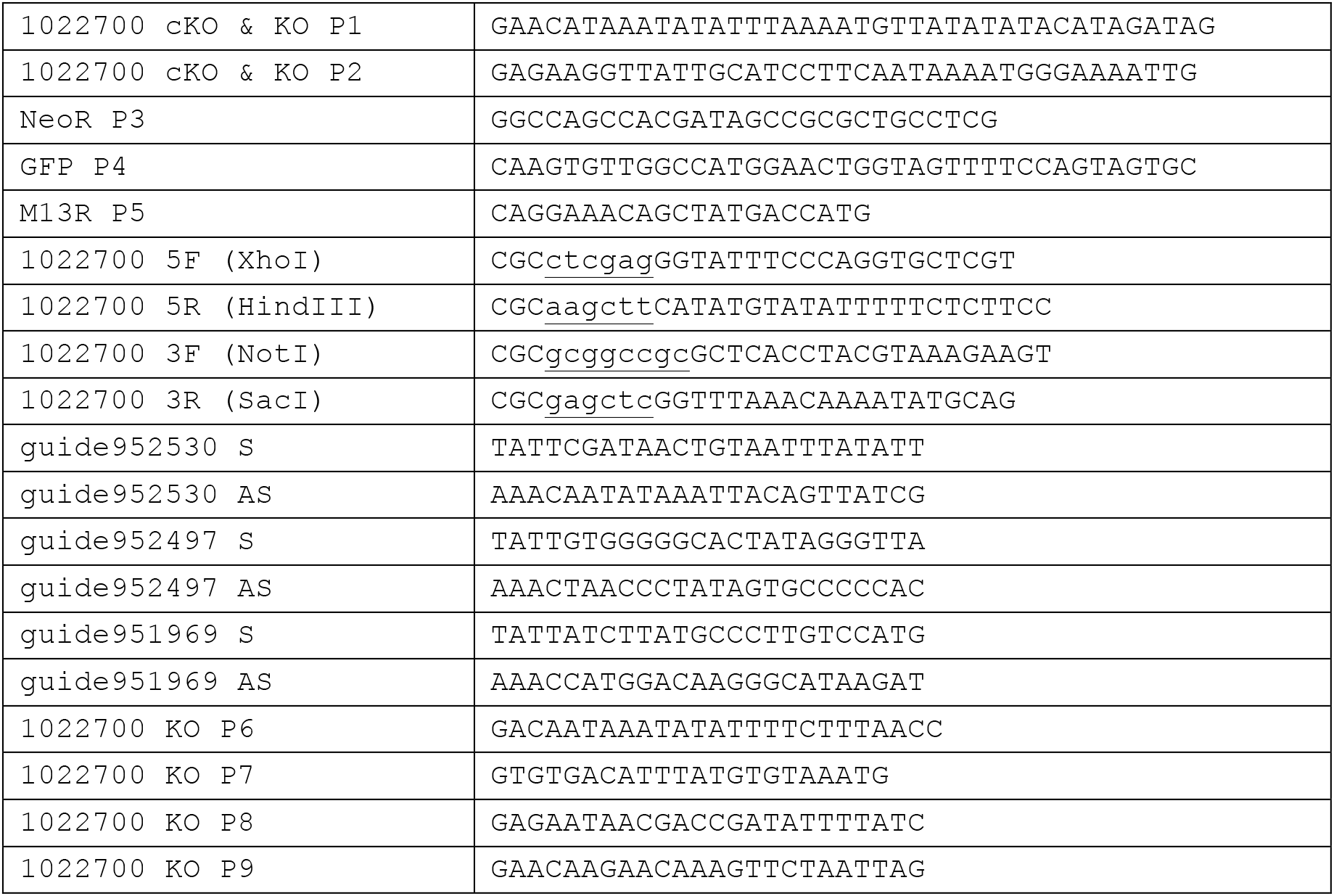
All primers used in the study.Table S1. All primers used in the study.

